# Evidence for a cohesion-related function of alpaca hums: Linking automated call detection with GPS data

**DOI:** 10.64898/2026.10.11.758387

**Authors:** Daniel Ariza-Echeverri, Riley O. Hayes Revitt, Stephan T. Leu, Kaja Wierucka

## Abstract

Understanding how animals communicate while moving helps explain social dynamics, but simultaneously tracking vocal and spatial behaviour in freely moving groups remains challenging. Animal-borne audio and GPS recordings can place naturally occurring vocalisations in their spatial context, but continuous recordings generate large datasets that require automated call detection. We used this approach to investigate whether alpaca hums, a poorly characterised vocalisation, contribute to group cohesion. We combined automated detection of hums from continuous collar-borne audio with synchronised GPS data from a partially collared herd to test whether humming is associated with changes in spacing and movement consistent with a contact-call function. We combined deep-learning-assisted detection with manual validation to build a large curated hum corpus from 2,011.1 h of continuous recordings from a partially collared herd. We then tested how humming was associated with spacing and movement at both individual and group levels. Following hum production, distance between focal alpaca and the group shifted toward closer contact relative to matched non-hum periods. Hums were also more likely to occur when the collared alpaca moved faster. These findings support a cohesion-related role of hums and suggest that this call type may help alpaca maintain social contact while moving. More broadly, our study shows how automated call detection combined with movement data can place poorly characterised vocalisations in a behavioural context, narrow their plausible functions, and generate targeted hypotheses even when caller identity remains uncertain and only part of a group can be collared. This approach extends functional research on vocal communication to species whose calls remain poorly understood.

## INTRODUCTION

Understanding how animals behave and communicate is the basis for explaining how social groups form and adapt to environmental changes (Demartsev et al. 2023). In group-living species, communication and collective movement depend on the social and spatial relationships among individuals, including who is nearby, how they move and how their spatial arrangement changes (Xie et al. 2024), but also on the relationships among individuals as they move and communicate (Demartsev et al. 2024). Investigating behaviour in multiple animals simultaneously is difficult, as complexity increases quickly with group size. Continuously tracking simultaneous movement and vocal activity of several freely moving animals is often not possible for a human observer. Sustained observation also requires substantial time, and is susceptible to visibility limits and bias (Demartsev et al. 2023; Oestreich et al. 2024; Enari et al. 2026). Biologging offers an alternative approach in which animal-mounted devices collect time-referenced measurements. Depending on sensor types, this can include location, acceleration, sound, physiological measures and environmental variables (Rutz and Hays 2009). Such devices can collect measurements repeatedly without a human observer and, when several animals are equipped, provide concurrent data streams across individuals (Johnson et al. 2009; Greif et al. 2019).

For studies on vocal communication, combining acoustic and movement sensors allows calls to be interpreted in their spatial and behavioural context (Demartsev et al. 2023). For example, animal-borne microphones capture the vocalisations of a collared animal, while satellite positioning records time-referenced trajectories. Synchronising the two allows each vocal event to be related to movement before, during and after it (Johnson and Tyack 2003; Casoli et al. 2022; Demartsev et al. 2023). Paired audio and location data have linked sound to movement across taxa, from marine mammals to social foraging in bats, revealing feeding bouts and nearby animals that location data alone could miss (Johnson and Tyack 2003; Egert-Berg et al. 2018; Greif et al. 2019; Oestreich et al. 2024). Recent terrestrial studies have extended this approach to free-ranging social mammals. In meerkats, simultaneous GPS and collar-borne audio recordings from multiple group members have been used to reconstruct the spatial and temporal structure of vocal interactions, showing that close-call exchanges vary with the animals’ local social environment (Demartsev et al. 2024). Multi-sensor tracking of white-nosed coatis has linked vocal activity and movement to fission-fusion events and collective travel decisions (Grout et al. 2026; Winans et al. 2026). Related multisensor deployments in spotted hyenas have enabled fine-scale reconstruction of fission-fusion dynamics from simultaneous GPS and accelerometry, although the acoustic data collected by the collars were not analysed in that study (Strauss et al. 2024). In a complementary application, Enari et al. (2026) combined collar-borne acoustic recording with GPS to quantify context-dependent variation in deer rutting-call rates and evaluate its implications for passive acoustic monitoring.

While on-animal microphones allow acoustic recordings to be combined with concurrent movement data, they generate large volumes of continuous audio that must first be processed, making the extraction of biologically relevant information a major bottleneck. Often, vocal events are detected by manually inspecting recordings or spectrograms, an approach that becomes increasingly impractical as recording duration and the number of simultaneously recorded animals increase (Demartsev et al. 2023; Kershenbaum et al. 2025). Automated call detection addresses this problem by computationally scanning recordings and identifying segments that are likely to contain a target vocalisation, allowing human effort to be concentrated on labelling training examples and validation rather than exhaustive review of the complete archive (Knight et al. 2020; Webber et al. 2022; Kershenbaum et al. 2025). Earlier automated workflows used hidden Markov models, signal-based detectors and machine-learning classifiers, often with operator validation (Knight et al. 2020; Webber et al. 2022). More recently, deep-learning frameworks have learned target sounds from labelled examples and applied the resulting detectors across much larger quantities of audio (Bergler et al. 2022). This allows vocal behaviour to be analysed across much larger quantities of audio recordings. Calls can be extracted from continuous recordings and linked to concurrent movement data and group configuration rather than being restricted to recordings that can feasibly be annotated by hand (Demartsev et al. 2023). This is particularly useful for species with poorly characterised vocalisations, because large numbers of naturally occurring calls can be recorded, extracted and related to their behavioural context, allowing to understand the calls (Oestreich et al. 2024). However, automated segmentation tools are often developed for specific species. Examples include detectors for mice, marmosets, killer whales, and elephants (Van Segbroeck et al. 2017; Zhang et al. 2018; Bergler et al. 2019; Zeppelzauer et al. 2015), which may not generalise reliably to other species without retraining or adaptation (Kershenbaum et al. 2025). This challenge may be compounded in mammals, whose vocalisations often vary gradually rather than falling into discrete call types, making individual call types more difficult to distinguish (Wadewitz et al. 2015; Kershenbaum et al. 2025).

The combination of large-scale call extraction and synchronised movement data is particularly well suited to investigating how vocalisations contribute to group cohesion. When animals forage or travel together, often without continuous visual contact, vocalisations can provide information about the presence and identity of nearby group members (Xie et al. 2024). Contact calls occur across diverse avian and mammalian social systems and can convey information about the caller and have also been associated with changes in group spacing and cohesion, helping groups remain together or enabling separated individuals to rejoin (Kondo and Watanabe 2009). However, how vocalisations influence group dynamics is only partly understood (Xie et al. 2024). Empirical studies provide specific examples of these links: movement calls recruit followers in dwarf mongooses (Cobb et al. 2022), vocal communication contributes to spatial coordination during flocking in zebra finches (Arnold et al. 2022), and increased grunt rates accompany collective departures in red-fronted lemurs, consistent with a role in coordinating group movement (Sperber et al. 2017). However, the terminology used across studies remains inconsistent, which complicates comparing such findings across different species. Terms such as close call, isolation call, lost call and contact call blend several criteria: the caller’s situation, the call’s presumed range, the behaviour it accompanies and its supposed outcome. Quiet short-range calls among foragers are often described as close calls because of how they sound and the context in which they are produced, rather than a specific communicative function (Jansen et al. 2012; Gall and Manser 2017; Reber et al. 2013).

Establishing call function usually requires several complementary forms of evidence (Oestreich et al. 2024). Bioacoustic analysis tests whether calls encode specific information, but not whether receivers use it. Playback experiments test receiver responses under controlled conditions, but they alter the natural context in which calls are produced and received (Berg et al. 2011; Sharpe et al. 2013; Rubow et al. 2017). Synchronised audio and movement recordings complement these approaches by relating naturally occurring calls to how collared animals are spatially arranged relative to each other and how they move before and after a call occurred (Demartsev et al. 2023; Casoli et al. 2022). For example, vocal activity has been related to subsequent changes in movement or spacing across several social species. In meerkats, moving-call choruses precede increases in group speed and cohesion-related changes between foraging patches (Bousquet et al. 2011), while close-call production varies with the spatial arrangement of group members and is linked to group cohesion during foraging (Engesser and Manser 2022). In elephants, rumbles have been shown to be followed by reduced interpartner distances (Leighty et al. 2008), while contact calls have shown associations with group contraction in goats (O’Bryan et al. 2019). In such cases, these analyses built on prior knowledge about the study species and its vocal repertoire. Applying the same approach to a less well-characterised vocal system offers a complementary opportunity. Large datasets containing simultaneous acoustic, spatial and behavioural observations can reveal consistent patterns even when baseline knowledge is limited, thereby supporting the development of testable hypotheses about call function. Alpaca provide a suitable study system for demonstrating this approach.

Alpaca are domesticated camelids native to South America (Pilarczyk et al. 2025). They are now raised in many countries worldwide and have become increasingly popular in Europe, where they are kept for fibre production, breeding and recreation, including on small, often hobby-oriented holdings (Neubert et al. 2021; Ząbek et al. 2024; Pilarczyk et al. 2025). Despite this growing popularity, scientific knowledge of alpaca behaviour remains scarce and dispersed (Miranda-de la Lama and Villarroel 2023). Even basic aspects of their social organisation remain poorly characterised. Alpaca are consistently described as gregarious and herd-living, but it remains unclear how persistent their groups are or how often individuals split from and rejoin them (fission-fusion events). In contrast, group dynamics have been characterised in other social mammals, including stable territorial groups in meerkats (Duncan et al. 2021) and fission–fusion clans in spotted hyenas (Smith et al. 2008). Available evidence nevertheless shows that social contact is important to alpaca: experimental spatial and visual isolation from familiar conspecifics increased heart rate and movement and reduced feeding compared with the presence of familiar companions (Pollard and Littlejohn 1995). How alpaca maintain contact during normal group activity remains unclear.

Alpaca vocal behaviour is similarly poorly characterised. Although several vocalisations have been described (Fowler 2008; Miranda-de la Lama and Villarroel 2023), their vocal repertoire has not been systematically characterised, unlike the well-described repertoires of species such as meerkats and African elephants (Manser et al. 2014; Soltis 2010). Hums are described as the most common alpaca vocalisation (Fowler 2008; Miranda-de la Lama and Villarroel 2023). They has a low and largely flat contour and encodes caller identity (Wierucka and Leu 2024). However, when hums are produced during naturally occurring social behaviours and what function they serve remain poorly understood. Alpaca spend much of their daytime activity grazing, during which feeding is accompanied by locomotion while searching for food (Ząbek et al. 2024; Boughey et al. 2025). If hums contribute to maintaining social contact, their production should therefore be related to how animals move and change their spacing during this routine activity.

Here, we investigated when hums occur in relation to group movement and spacing, and whether spatial changes following hums are consistent with a cohesion-related function. We developed a pipeline for automatically detecting alpaca vocalisations from continuous recordings. We then related the detected hums to synchronised movement data and tested whether humming was associated with spatial patterns predicted by two candidate cohesion-related functions. We approached this at two complementary levels: First, at the individual-level, we asked whether an animal’s distance to its group changes after it calls. Second, at the group-level, we asked in which spatial states calling is most frequent. Based on this, we tested two complementary hypotheses (H). First, we hypothesised that humming restores or reinforces contact when an alpaca is separated from the group (H1 Contact Reinforcement), and investigated what happens after a hum. This predicts that the caller’s distance to the group centroid decreases following a hum. Second, at the group level, we hypothesised that hums help maintain contact while group members reposition (H2 Contact Maintenance). Here, we investigated whether hums occur when group spacing changes. This predicts that humming is more frequent when the group begins to spread while its members are still relatively close together.

## MATERIALS AND METHODS

### Data collection

#### Study site and data collection

We collected data at Adelaide University’s Rose-worthy campus, South Australia, in February 2025, across 11 daily recording sessions. The fenced paddock covered approximately 0.7 ha and contained shade, hay and fresh water. We fitted c ollars to 15 of approximately 50 alpaca in the herd and released the animals into the paddock. The 15 collars recorded concurrently during the daily recording sessions. The University of Adelaide Animal Ethics Committee approved all procedures (approval number S-2024-102).

Each collar carried an AudioMoth Dev recorder with an AudioMoth GPS Board (Open Acoustic Devices) in a custom-made protective case. Audio was recorded at 48 kHz with gain set to 3, and GPS positions were recorded at a nominal sampling rate of 1 Hz. Recordings ran from 06:00 to 21:00 Australian Central Daylight Time (ACDT; UTC+10:30). GPS and audio data were synchronised automatically using a pulse-per-second signal embedded in the GPS module (one microsecond accuracy).

### Constructing the hum corpus

#### Recordings and Manual Annotation

*Hums.* We defined a hum following existing definitions as a vocalisation with a predominantly flat peak-frequency contour, allowing brief frequency modulation at the beginning or end of the call (Wierucka and Leu 2024; Fig. 1). Previous analyses have characterised hums with a mean Fundamental Frequency (*F*_0_) of 272 Hz, mean Peak Frequency of 353 Hz, and a mean Duration 90% of 464 ms. Mean Frequency 5% and Frequency 95% were 251 and 828 Hz, respectively, corresponding to a mean Bandwidth 90% of 577 Hz (Wierucka and Leu 2024).

**Figure 1.**
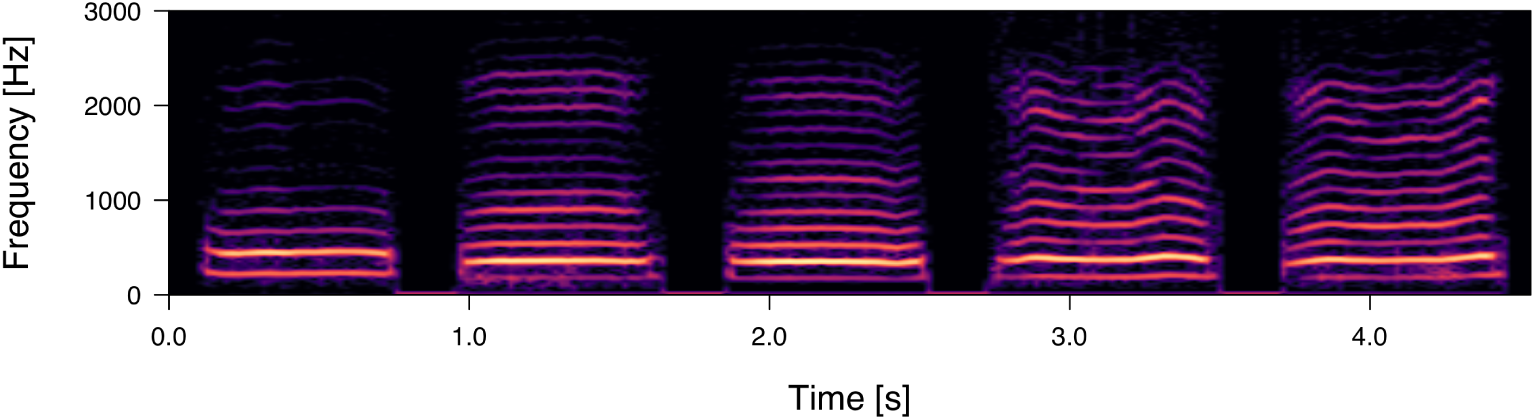
Example spectrograms of alpaca hums. Hums were identified by a comparatively flat frequency contour and harmonic structure; brief frequency modulation could occur near call onset or offset. Spectrogram settings: DFT size 2,048 samples, Hann window 41.7 ms, hop size 20.8 ms, with 50% overlap.

To create a labelled dataset for training and evaluating the model for automated hum detection, we manually annotated a subset of the recordings. This subset consisted of approximately 1 h recording segments from multiple days. We used Raven Pro (ver. 1.6.5), and all hums, as defined above, were identified through visual and auditory inspection of the spectrograms. No hums were excluded based on recording quality.

#### Automated Detection of Alpaca Hums

To generate candidate hums from the continuous recordings, we used ANIMAL-SPOT (Bergler et al. 2022) as the basis for a deep learning vocalisation detection workflow, comprising training data preparation, model training and sliding window detection. We retained its classifier architecture and adapted the parts of the workflow pertaining to data ingestion, sampling of noise excerpts and added downstream post-filtering layers. Additionally, we tuned its input and inference settings to the acoustic profile of alpaca hums.

To create a training dataset containing both hum and noise excerpts, we added a preprocessing step to ANIMAL-SPOT that sampled noise from sections of the annotated recordings outside the identified hum intervals. We used these excerpts to expose the detector to diverse background sounds while retaining enough hum examples, prioritising high recall during candidate generation (Bergler et al. 2022; Knight et al. 2020). Each sampled noise excerpt matched the duration of a hum example, and sampling avoided the 0.5s immediately before and after each annotated hum to reduce call leakage. We subsampled the noise excerpts to a 1.5:1 noise-to-hum ratio to limit class imbalance while retaining diverse background examples (Bergler et al. 2022). We also manually verified sampled noise excerpts before model training. We allocated excerpts to training, validation and test partitions in a target ratio of 70:15:15.

We used a convolutional neural network (CNN), which analyses short spectrogram excerpts as image-like time—frequency inputs. ANIMAL-SPOT uses a ResNet-18 CNN architecture, an approach validated by the framework for target versus noise detection across multiple animal datasets (Bergler et al. 2022; He et al. 2016). We extracted 400-ms excerpts from the recordings and represented each as a linear frequency spectrogram with 128 bins between 0 and 4,000 Hz, using a 2,048 sample Fourier transform and a 1,024 sample hop. We trained for 40 epochs with a batch size of 16 and the Adam optimiser at a learning rate of 10*-*5 (Kingma and Ba 2015), keeping the checkpoint with the highest validation accuracy. We then applied the detector to all recordings, moving a 300-ms window through each, in 50-ms steps. The detector assigned each segment a model score, with higher values indicating stronger support for the target class. We manually inspected model predictions and found that retaining segments with scores of 0.5 and higher balanced candidate coverage against precision, and we merged temporally overlapping retained segments into candidate detections.

#### Candidate Curation and Duplicate Removal

The detector generated a broad candidate set that included false detections. We extended the ANIMAL-SPOT workflow, adding two supervised curation layers. This follows detector plus review approaches in which automation first narrows a large archive to candidate events and human labelled candidates are then used to remove systematic false positives before ecological analysis (Knight et al. 2020; Webber et al. 2022).

For the first layer we manually labelled candidates, classifying them as hums or noise. The noise class included low-amplitude, predominantly low-frequency sounds lacking a clearly identifiable hum structure, which we labelled *rumbles*, as well as hum-like sounds that were too faint to retain. We excluded these faint detections to reduce the inclusion of calls that may have originated from uncollared alpaca. Using Python 3.11 and librosa 0.11.0 (McFee et al. 2015), we extracted mel-frequency cepstral coefficients and spectro-temporal summaries of frequency, energy and duration. These feature classes have been shown to encode individual identity in alpaca hums, and random forests have successfully classified alpaca identity from those acoustic features (Wierucka and Leu 2024). We therefore used a random-forest classifier based on these features (scikit-learn 1.6.1; Pedregosa et al. 2011) for this first curation layer. We fitted it on a grouped 75:25 partition and selected the configuration with the highest F1 score from 150 model and decision threshold combinations. We retained candidates with a target class probability score of at least 0.60.

Manual inspection showed recurrent contamination by human speech and low-frequency noise, so we added a second layer of curation. We used class balanced logistic regression as a simple binary classifier for this narrower contamination removal step (Maalouf and Siddiqi 2014). Class balancing reduced the influence of the unequal hum and contaminant sample sizes. We fitted the model to manually reviewed detections labelled as hums or contaminants, using 34 spectro-temporal features and cepstral coefficients informed by identity encoding reported for alpaca hums (Wierucka and Leu 2024). We evaluated classifier performance using five-fold cross-validation. After evaluation, we applied the classifier to the entire dataset and retained detections with target class probability scores of at least 0.5. We chose this probability threshold to avoid imposing an additional strict cutoff that could disproportionately reduce recall, given that the candidate set had already undergone substantial curation by the preceding random forest classifier.

Because the same hum could be recorded by several nearby collars, we also removed repeated recordings of the same occurrence before spatial analysis. We identified plausible duplicates from their temporal and spatial proximity and then required acoustic agreement across seven frequency-, duration– and entropy-based descriptors. We evaluated 500 combinations of the temporal, spatial and acoustic thresholds and selected the configuration on the basis of acoustic coherence and stability of duplicate decisions and removal counts rather than maximum removal (Supplementary Materials (SM), Table S1). Because a call recorded on the caller’s own collar is expected to have a comparatively strong and clear acoustic signal, prioritising these characteristics also increased the likelihood that the retained recording originated from the caller’s collar. Therefore, one representative was retained from each duplicate group, prioritising signal to noise ratio, root mean square amplitude and detector score. For subsequent statistical analyses, we thus assumed that the retained recording was most likely to have originated from the caller’s own collar, characterized by being the loudest among possible duplicates.

### Spatial and movement measures

We assigned each GPS fix to the nearest second. Although GPS positions were recorded at a nominal rate of 1 Hz, these timestamps contained subsecond offsets, so rounding occasionally assigned two consecutive fixes from the same collar to the same second; in these cases, we used the median latitude and longitude. We then projected coordinates into a common local metric system. Missing GPS fixes were retained as gaps; positions were not interpolated.

GPS locations are subject to positional error that varies with environmental conditions, sampling context and device performance (Frair et al. 2010; Acácio et al. 2022). We assumed this error to be approximately normally distributed around true positions and unrelated to hum occurrence. As a quality-control step, we excluded GPS fixes located outside the study paddock, as these represented locations that were not physically accessible to the animals and were therefore considered erroneous. To represent the observed group’s spatial arrangement, we used complementary measures of group centre (the centroid) and group spread. We assumed that the collared animals were approximately randomly distributed throughout the herd, such that, on average, the centroid of the non-focal collared animals provided a representative estimate of the group’s spatial centre. First, we calculated the distance between a focal individual’s location and the group’s centroid (we excluded the focal animal’s position in the group centroid) as a reference for the individual’s position relative to the group. This measure has been used before in audio–GPS studies of contact calling (O’Bryan et al. 2019). Second, we calculated the convex-hull area as the two-dimensional area occupied by the group (Romenskyy et al. 2017). At each second, we therefore defined the group centroid as the arithmetic mean of the available positions and group spread as the area of the smallest convex polygon enclosing all collar positions. Missing GPS positions were retained as gaps and were not interpolated. Spatial measures were calculated only for seconds in which positions were available for at least 8 of the 15 collared alpaca, requiring GPS coverage from more than half of the collared animals; otherwise, the corresponding measure was treated as missing for that second.

### Statistical analyses

#### Hypothesis 1: Contact Reinforcement

We modelled H1 to test whether a hum marks a change in an individual alpaca’s spatial relationship to the observed group: specifically, whether focal-to-group distance changes differently after hum production than after comparable non-hum moments. Matched controls have been used to provide a baseline for how behaviour changes under similar conditions when the focal event (the hum) does not occur (Voelkl et al. 2016).

*Hum occurrences and distance change.* We defined an occurrence as a focal time point with a specified temporal and spatial context. A hum occurrence was a hum detected on the focal alpaca’s collar, whereas a control occurrence was a matched time point with no detected hum. For each hum occurrence, the occurrence time *t*_0_ was the midpoint between its start and end timestamps. We kept the exact midpoint time of each hum, while GPS positions from different collars were aligned to common one-second time points. At each GPS second we calculated the Euclidean distance between the focal animal and the centroid of the others. For each occurrence, we defined equally long analysis intervals before and after *t*_0_. For an interval length *w*, the pre-occurrence interval was [*t*_0_*−w, t*_0_) and the post-occurrence interval was (*t*_0_*, t*_0_+*w*]. We summarised focal-to-centroid distance by its median within each interval and calculated

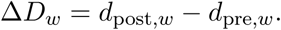

For each call occurrence, Δ*D_w_* therefore describes the change in focal-to-centroid distance from before to after that occurrence. A negative value means that the focal alpaca was closer to the centroid afterwards, whereas a positive value means that it was farther away. We calculated Δ*D_w_* in the same way for hum and control occurrences.

*Matched controls.* We selected non-hum control moments from the same animal and recording session and with the same interval length as for each hum occurrence. Controls had to come from the same collar, occur within 60 min of the hum detection, and differ by no more than 0.5 standard deviations in both *starting focal-to-centroid distance* and *pre-occurrence mean movement speed* of the focal animal. Candidate controls whose full analysis interval included hums were excluded.

*Linear models.* We fitted Gaussian linear mixed-effects models with Δ*D_w_* as the response variable. We expressed Δ*D_w_* in centimetres to guarantee model fit stability against potentially small decimal numbers, but we did not assume this level of GPS precision. Because an immediate movement adjustment in response to the hum and a broader reunion response may act on different time scales, we used 60-s intervals (Δ*D*_60_) for the primary 60-s H1 analysis and repeated the same analysis at 30s and 120s. The predictor of interest was *hum presence*. We denote its coefficient as *β*_hum_. A negative *β*_hum_ means that Δ*D_w_* was on average lower after hum production than after controls. This can reflect a stronger contraction, i.e. decrease in focal-to-centroid distance after hum production. But it can also reflect a weaker expansion, i.e. increase in distance after hum production. Thirdly, it can reflect a contraction after hum production, and an expansion in the control. All three scenarios, represented by a negative *β*_hum_ suggest a stronger social cohesion following hums relative to the control. Conversely, a positive *β*_hum_ means that Δ*D_w_* was on average larger after hum production than after controls. A negative *β*_hum_ would therefore provide evidence supporting H1 (Contact Reinforcement). A *β*_hum_ near zero would indicate no detectable difference between hum and control occurrences, whereas a positive coefficient would be inconsistent with the predicted cohesion effect (Supplementary Materials; Table S2).

To determine whether lower Δ*D*_60_ after hum production than after matched controls mainly reflected decreases in focal-to-centroid distance, we examined pairs in which Δ*D*_60_ was lower after the hum. We distinguished two cases: distance decreased after the hum (combining the first and third scenarios described above), or distance increased in both conditions but less after the hum. We then calculated how much each case contributed to the lower Δ*D*_60_ after hum production than after matched controls.

To ensure that *β*_hum_ did not simply capture an existing tendency toward greater or lower cohesion, we also included five measures of spatial and movement state before the hum or control as fixed effects: the *starting focal-to-centroid distance* (*d*_pre_*_,w_*), representing how far the focal alpaca initially was from the group; pre-occurrence (*t*_0_ – *w, t*_0_) mean focal speed, representing its movement activity; *recent focal-to-centroid distance change*, calculated as *d*_pre_*_,w_* minus the median focal-to-centroid distance in the immediately preceding interval [*t*_0_ – *2w, t*_0_ – *w*), indicating whether focal-to-centroid distance was already decreasing or increasing; median group spread within the pre-occurrence interval (*t*_0_ *– w, t*_0_), representing how dispersed the group was; and *recent group spread change* was this value minus the median group spread in the immediately preceding interval (*t*_0_ *– 2w, t*_0_ *– w*), indicating whether the group was already expanding or contracting. We log-transformed *starting focal-to-centroid distance* and *pre-occurrence mean focal speed*, which were right skewed and bounded at zero, and standardised all continuous predictors to units of one standard deviation. The 30-s and 120-s models used the same predictor structure with their corresponding interval measures.

To account for repeated observations, we included random intercepts for focal alpaca identity and recording session. To avoid overconfident fixed-effect estimates, we also included all theoretically identifiable random slopes (Schielzeth and Forstmeier 2009; Barr et al. 2013). Specifically, we fitted random slopes for *hum presence* and each of the five continuous predictors within both focal alpaca identity and recording session, and estimated correlations among the corresponding random intercepts and slopes.

We fitted all statistical models in R 4.3.1 (R Core Team 2023; lme4 2.0-6 package (Bates et al. 2015)). We tested *β*_hum_ by comparing the full model with a null model omitting that term, using maximum-likelihood fits and a likelihood-ratio test. Additionally, we assessed the contribution of the five spatial and movement state fixed effects by comparing maximum-likelihood fits of the full model with models omitting each fixed effect in turn by computing AIC. These comparisons favoured the full fixed-effect set for every time interval. To assess the stability of *β*_hum_ in the primary 60-s H1 analysis, we refitted the model after excluding each focal alpaca and each recording session in turn. We also assessed collinearity among the fixed-effect predictors in each H1 model using variance inflation factors (VIFs); maximum VIFs were 1.40, 1.49 and 1.66 for the 30-s, 60-s and 120-s analyses, respectively.

*Contributions to distance change and sensitivity analysis.* Because Δ*D_w_* could result from movement of the focal alpaca, movement of the group centroid, or both, we examined these movements separately. For each occurrence, we calculated the median positions of the focal alpaca and the centroid in the pre– and post-occurrence intervals. To assess the focal alpaca’s movement, we held the centroid at its pre-occurrence position and compared the focal alpaca’s distance to that fixed point before and after the occurrence. A decrease in this distance indicated that the focal alpaca had moved closer to the centroid’s earlier position. We then repeated the comparison for centroid movement, this time holding the focal alpaca at its pre-occurrence position and comparing its distance to the centroid before and after the occurrence. A decrease indicated that the centroid had moved closer to the focal alpaca’s earlier position. We used these two comparisons to distinguish whether the focal alpaca, the centroid, or both moved toward the other’s earlier position.

*Sensitivity to distant alpaca.* Because unusually distant collared alpaca could disproportionately influence the group centroid, we repeated the H1 analysis after excluding their positions beyond a distance threshold. To determine this threshold, we calculated the median distance between each non-focal collared alpaca and the group centroid during the 60-s pre-occurrence interval. Unlike the *starting focal-to-centroid distance* used in the primary H1 analysis, which measured the focal alpaca’s distance to the centroid, these distances were calculated for each non-focal collared alpaca. We pooled these median distances and defined the threshold using Tukey’s upper fence (Q3 + 1.5 *×* IQR; Tukey 1977).

#### Hypothesis 2: Contact Maintenance

We modelled H2 to test whether hum production depended on the observed group’s spatial state, specifically whether hums were more frequent when a compact group was beginning to spread. Here, we shifted from the individual level (hum–control) comparison in H1 to a group-level analysis, using the number of hum occurrences in each 60-s time window as the response and relating it to measures calculated across the collared group.

*Dataset preparation.* Because not all alpaca were collared, a collar-mounted AudioMoth could capture a hum produced by an uncollared animal. Such calls might not be distributed evenly across group states: when the collared animals were close together, uncollared neighbours could also be nearby. Hence it is more likely that time windows of low group spread could include hums that were not produced by the collared alpaca, relative to time windows of larger spread.

Because the AudioMoths were mounted on the animal, we assumed that hums produced by the focal animal (wearing the collar) would generally be of higher amplitude than hums produced by nearby animals. We therefore used the amplitude of calls in recordings to remove quieter hums that were more likely to have come from another, uncollared alpaca. A similar amplitude-based approach has been used with synchronised animal-borne tags to distinguish focal from non-focal calls (Zeh et al. 2024). We calibrated amplitude thresholds via a separate controlled recording experiment. Two focal alpaca were fitted with collars and observed in separate sessions, while four stationary AudioMoths were positioned 1, 2, 4 and 8 m from the caller. Across the two sessions, the focal alpaca produced 212 and 162 hums, respectively (374 in total). For each hum, we measured its amplitude on the caller’s own collar and on each of the four stationary AudioMoths. We pooled these stationary AudioMoth measurements and used the 90th, 95th and 97.5th percentiles of their amplitude distribution as candidate thresholds (*-*35.75, *-*33.06 and *-*30.85 dBFS, respectively; Supplementary Fig. S1).

At the 95th-percentile threshold of *-*33.06 dBFS, 94.6% of hums recorded on the caller’s own collar were retained, whereas only 5.0% of the corresponding recordings on the stationary AudioMoths reached or exceeded the threshold. Based on this calibration, we used *-*33.06 dBFS to construct the dataset for the H2 statistical analyses, retaining hum detections at or above this amplitude and excluding quieter detections. We retained the 90th and 97.5th percentile thresholds to further test how sensitive the results were to the percentile choice.

For H1, we retained all curated hum detections because the response variable, Δ*D_W_*, was calculated from GPS positions, while acoustic detections determined the occurrence times and focal animal assignments. In contrast, H2 used *hum count* as the response, which could be directly inflated by calls from nearby uncollared alpaca, particularly when the group was compact. Because this source of error could vary with *group spread* being low, it could systematically amplify the bias between the response variable and one of the main predictors of interest (Keogh et al. 2020). We therefore applied the additional amplitude threshold only to H2.

*Hum counts and group measures.* To pair hum production with a concurrent group state, we divided each recording session into non-overlapping 60-s time windows and counted hum occurrences retained above the 95th percentile amplitude threshold, whose midpoint fell within each window; incomplete terminal windows were discarded. We considered 30-s and 120-s windows before selecting 60s. Compared with 30s, a one-minute window provided more temporal margin to capture humming that may occur in bouts, whereas compared with 120s it retained a larger number of non-overlapping observations for analysis. We used 60-s time windows throughout H2, so hum counts and group state predictors referred to the same one-minute period.

*Current spread* was the 60-s group spread summary defined in the spatial and movement measures subsection above, and *spread change* was the corresponding difference from the preceding window, expressed per minute. *Mean group member speed* was calculated from the distance travelled by each collared alpaca between consecutive GPS positions recorded 1s apart, averaged across all collared alpaca within each window. We excluded speed estimates exceeding 12 m/s as biologically implausible; such exclusions have also been applied in previous studies (Demartsev et al. 2024). *Centroid displacement* was the Euclidean distance between the current and preceding window centroids, each the average of the per-second centroids. It measures shift of the group centre, not cohesion, travel direction or movement synchrony. We retained predictors in their original measurement units for descriptive summaries and standardised continuous predictors to one-SD units before fitting.

*Linear models.* We modelled hum counts with generalised linear mixed-effects models in R 4.3.1 using glmmTMB 1.1.14 (O’Hara and Kotze 2010; Harrison et al. 2018; Brooks et al. 2017). We first fitted Poisson models (Lindén and Mäntyniemi 2011) and checked model assumptions with DHARMa 0.5.0 (Hartig et al. 2026), using 1,000 simulated residual datasets per fitted model to assess dispersion, zero inflation, and within-session temporal autocorrelation. The Poisson models showed overdispersion and excess zeros, so we used negative-binomial type 1 (NB1) models for H2 (Lindén and Mäntyniemi 2011). In the NB1 models fitted t o h ums r etained a bove t he 95th percentile amplitude threshold, DHARMa tests did not indicate remaining overdispersion or excess zeros, although residual temporal autocorrelation remained detectable within most recording sessions.

*Current spread*, *spread change*, *mean group member speed* and *centroid displacement* were included as predictors. *Current spread* and *spread change* were modelled with an interaction term, which tested whether the association between *spread change* and humming depended on how spread out the group already was. To account for repeated observations within recording sessions, we included a random intercept for recording session and random slopes for *current spread*, *spread change*, *mean group member speed*, *centroid displacement* and the *current-spread × spread-change* interaction. Window length in minutes was included as an exposure offset.

We report model estimates as incidence-rate ratios (IRRs). An IRR is the exponentiated coefficient from a count model and expresses the relative change in an expected event rate associated with a change in a predictor (Halvorson et al. 2022). In the H2 statistical models, the event rate was the expected number of hums per minute. Because continuous predictors were standardised, each IRR describes how this expected hum count changes for a one-SD increase in the corresponding predictor, while the other model terms are held constant. An IRR of 1 indicates no change in the expected hum count, an IRR above 1 indicates an increase, and an IRR below 1 indicates a decrease.

The *current-spread × spread-change* interaction is the primary H2 term. Under H2, the number of hums would increase most strongly with increasing *spread-change* when the group is still compact (low *current spread*), which corresponds to a negative interaction coefficient, alongside a negative association with *current spread* and a positive association with *spread change*. *Mean group member speed* and *centroid displacement* were included as additive terms, corresponding respectively to movement activity among group members and displacement of the group centre. Evidence for H2 (Contact Maintenance) would therefore come from an interaction IRR below one, with negative *current-spread* and positive *spread-change* associations providing additional support (Supplementary Table S3).

## RESULTS

### Corpus construction

The 15 alpaca were collared simultaneously, yielding a total of 2,011.1 h of paired acoustic and GPS data across collars. The training dataset for the CNN-based detector contained 4,325 labelled excerpts: 1,730 target hums and 2,595 noise examples. Applying the CNN detector to the 274 recordings produced 58,512 candidates. The random forest classifier curation layer retained 29,387 candidates (50.22%), and the logistic regression classifier secondary curation layer retained 28,821 detections (98.07% of its input). Manual review (GPS informed) excluded seven additional speech detections, leaving 28,814 detections. Duplicate removal retained 28,561 hum occurrences, removing 253 detections (0.88%) that represented additional recordings of the same occurrences. Overall, 28,561 hum occurrences were retained for analysis, representing 48.8% of the initial 58,512 candidate detections (Table 1; Fig. 2). Retained hums had a median duration of 0.55s (IQR: 0.45–0.70s), and 93.9% were shorter than the 1-s GPS sampling interval.

**Figure 2.**
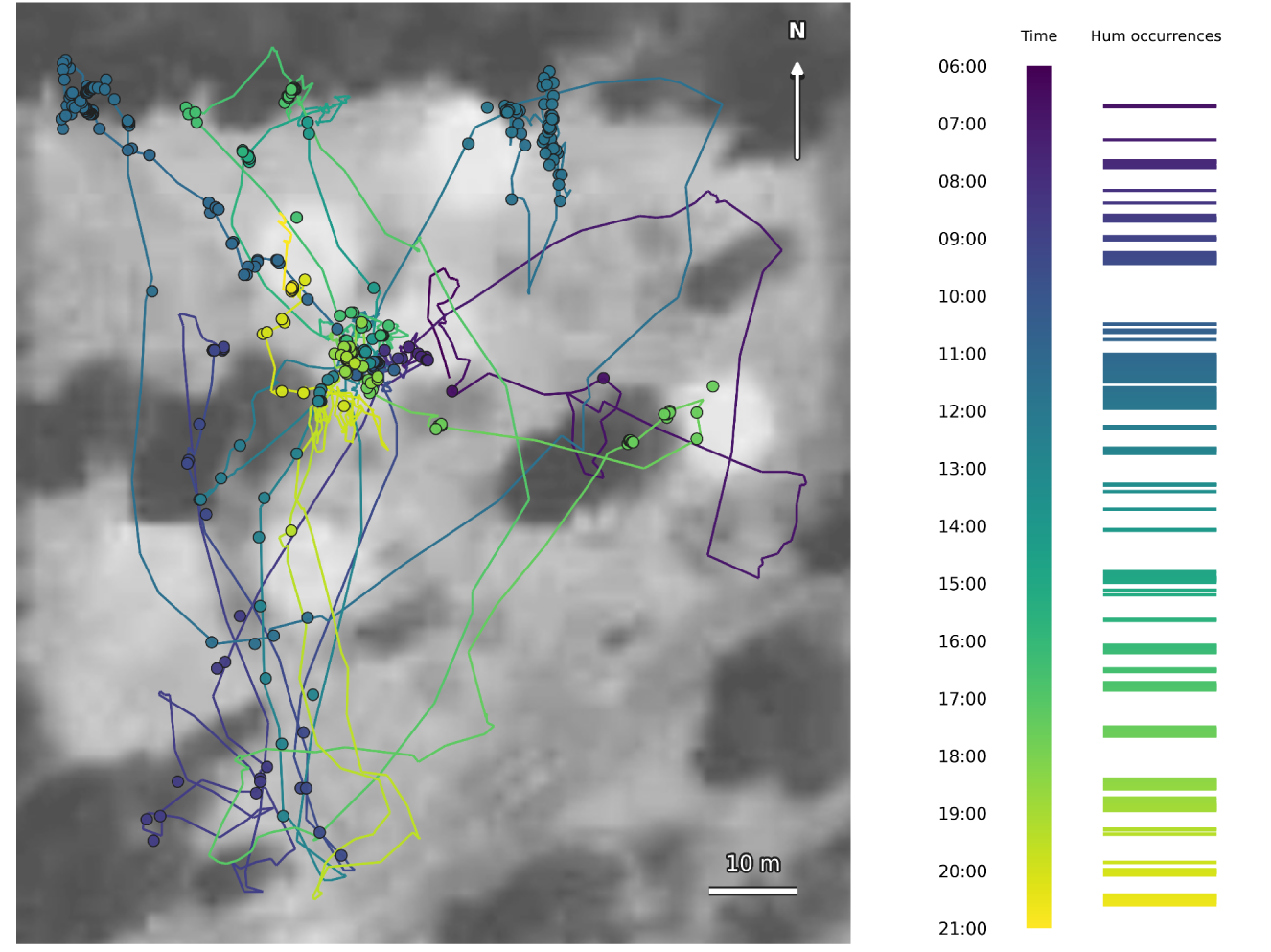
Representative example of movement and hum timing for a collared alpaca across one recording day. Colour represents time (local). Hum occurrences are marked next to the time scale and represented by coloured dots on the plot (n = 546; no amplitude threshold applied).

**Table 1.** Stagewise construction of the alpaca hum corpus. Percentages use the preceding layer as their denominator. The evaluation column reports the principal performance or robustness measures for each automated layer.

| Corpus layer | Input | Output | Change from input | Evaluation summary |
| --- | --- | --- | --- | --- |
| CNN candidate generation | 274 recordings (2,011.1 h) | 58,512 candidates | — | Test accuracy 0.968; F1 0.964; precision 0.970; recall 0.958 |
| Random-forest candidate curation | 58,512 candidates | 29,387 detections | 29,125 excluded; 50.22% retained | F1 0.749; precision 0.747; recall 0.752; ROC AUC 0.875 |
| Logistic contamination filtering | 29,387 detections | 28,821 detections | 566 excluded; 98.07% retained | Balanced accuracy 0.801; hum precision 0.935; hum recall 0.827; noise recall 0.775 |
| GPS-informed manual review | 28,821 detections | 28,814 detections | 7 confirmed speech detections excluded; 99.98% retained | Manual confirmation |
| Duplicate removal | 28,814 detections | 28,561 hums | 253 additional recordings removed; 99.12% retained | Local decision stability 0.958; removal-count stability 0.977 |

The selected CNN achieved an F1 score of 0.964 on the test set. The random forest achieved an F1 score of 0.749 on the test set, and the logistic contamination classifier achieved a five-fold cross-validated balanced ac curacy of 0. 801 (Supplementary Table S4).

### Hypothesis 1: Contact Reinforcement

In the primary 60-s H1 analysis, *β*_hum_ was negative and statistically supported (full– null likelihood-ratio test: *P* = 0.024; Table 2). *β*_hum_ was negative at all three interval lengths, meaning that Δ*D_w_* was on average lower when hums occured. *β*_hum_ became progressively lower with increasing interval length (Table 3). Among the other predictors, Δ*D*_60_ was lower when *starting focal-to-centroid distance* (*d*_pre_*_,w_*) was greater (P < 0.001), meaning that alpaca starting farther from the centroid tended to show a greater decrease, or smaller increase in distance. Δ*D_w_* was higher when *recent focal-to-centroid distance change* was greater (P < 0.001), meaning that distance tended to continue changing in the same direction, and when *pre-occurrence group spread* was greater (P = 0.007). The same three predictors were statistically supported at 30s. At 120s, only *d*_pre_*_,w_* remained statistically supported; *recent focal-to-centroid distance change* (P = 0.106) and *pre-occurrence group spread* (P = 0.139) were not. The effect of *pre-occurrence mean focal speed* and *recent group spread change* were not statistically significant at any interval.

**Table 2.** Fixed-effect estimates from the primary 60-s H1 (Contact Reinforcement) analysis, testing whether *focal-to-centroid distance change* (Δ*D*_60_) differed after hum production compared with matched non-hum control moments. The response was the difference in median distance between the focal alpaca and the centroid of the other collared alpaca in the 60s before and after each occurrence. The coefficient for hum presence (*β*_hum_) represents the adjusted difference between hum and control occurrences; negative *β*_hum_ indicates lower mean distance change after hum production. Estimates and 95% confidence intervals are expressed in centimetres. Continuous predictors were standardised to one standard deviation before model fitting. *P*-values are from coefficient-level Satterthwaite *t*-tests

| Predictor | Estimate [cm] | 95% CI [cm] | P |
| --- | --- | --- | --- |
| Intercept | 19.98 | -5.44 – 45.40 | 0.112 |
| Hum presence ($\beta_{\text{hum}}$) | -45.96 | -88.03 – -3.89 | 0.034 |
| Starting focal-to-centroid distance | -110.33 | -139.58 – -81.08 | < 0.001 |
| Pre-occurrence mean focal speed | 10.76 | -22.73 – 44.25 | 0.511 |
| Recent focal-to-centroid distance change | 81.38 | 53.98 – 108.77 | < 0.001 |
| Pre-occurrence group spread | 30.68 | 9.64 – 51.72 | 0.007 |
| Recent group spread change | 8.47 | -9.46 – 26.40 | 0.321 |

**Table 3.** Hum-presence *β*_hum_ estimates across H1 interval lengths. Estimates and 95% confidence intervals (CI) of focal-to-centroid distance change (Δ*D_w_*). A negative *β*_hum_ estimate indicates that mean Δ*D_w_* was lower when hums occurred than for matched non-hum controls. *P*_LRT_ values are from full–null likelihood-ratio tests fitted by maximum likelihood.

| Interval | Estimate [cm] | 95% CI [cm] | PLRT |
| --- | --- | --- | --- |
| 30s | -25.33 | -44.69 to -5.97 | 0.020 |
| 60s | -45.96 | -88.03 to -3.89 | 0.024 |
| 120s | -90.39 | -149.03 to -31.76 | 0.009 |

In the primary 60-s H1 analysis, the lower Δ*D*_60_ after hum production than matched controls was mainly driven by cases in which focal-to-centroid distance decreased after hum production (92.2%), rather than by cases in which distance increased in both hum and matched control conditions but increased less after hum production (7.8%; Supplementary Table S5). Before hums, focal-to-centroid distance was already decreasing in 51.8% of occurrences and increasing in 48.1%; these proportions were nearly identical at 30 and 120s.

After excluding positions of unusually distant collared alpaca, *β*_hum_ remained negative and statistically significant at all three interval lengths (30s: P = 0.014; 60s: P = 0.022; 120s: P = 0.007). The movement-source analysis showed that hum occurrences associated with increased cohesion involved movement of the focal alpaca toward the group centroid (36.1%), movement of both toward each other (34.3%), or movement of the centroid toward the focal alpaca (29.6%).

### Hypothesis 2: Contact Maintenance

The H2 analysis included 8,808 eligible 60-s time windows across 11 recording sessions, containing 4,617 hum occurrences retained above the 95th percentile amplitude threshold. Of these windows, 1,482 (16.8%) contained at least one retained hum and 7,326 (83.2%) contained no retained hum.

In the NB1 model, the full–null model comparison did not support the *current-spread × spread-change* interaction (likelihood-ratio test: P = 0.250); the interaction estimate was IRR: 0.965 (95% CI: 0.904–1.029). Thus, hums were not specifically more frequent when relatively compact groups were expanding, as predicted under H2. However, more hums were produced when the collared alpaca were moving faster (*mean group member speed*: IRR: 1.179, 95% CI: 1.119–1.243; P < 0.001; Table 4; Fig. 3). At the mean of the other interacting predictor, hum count did not vary detectably with *current spread* or *spread change*. *Centroid displacement* was also not statistically supported, meaning that hum production did not change detectably with how far the group centre shifted between consecutive 60-s windows (Table 4).

**Figure 3.**
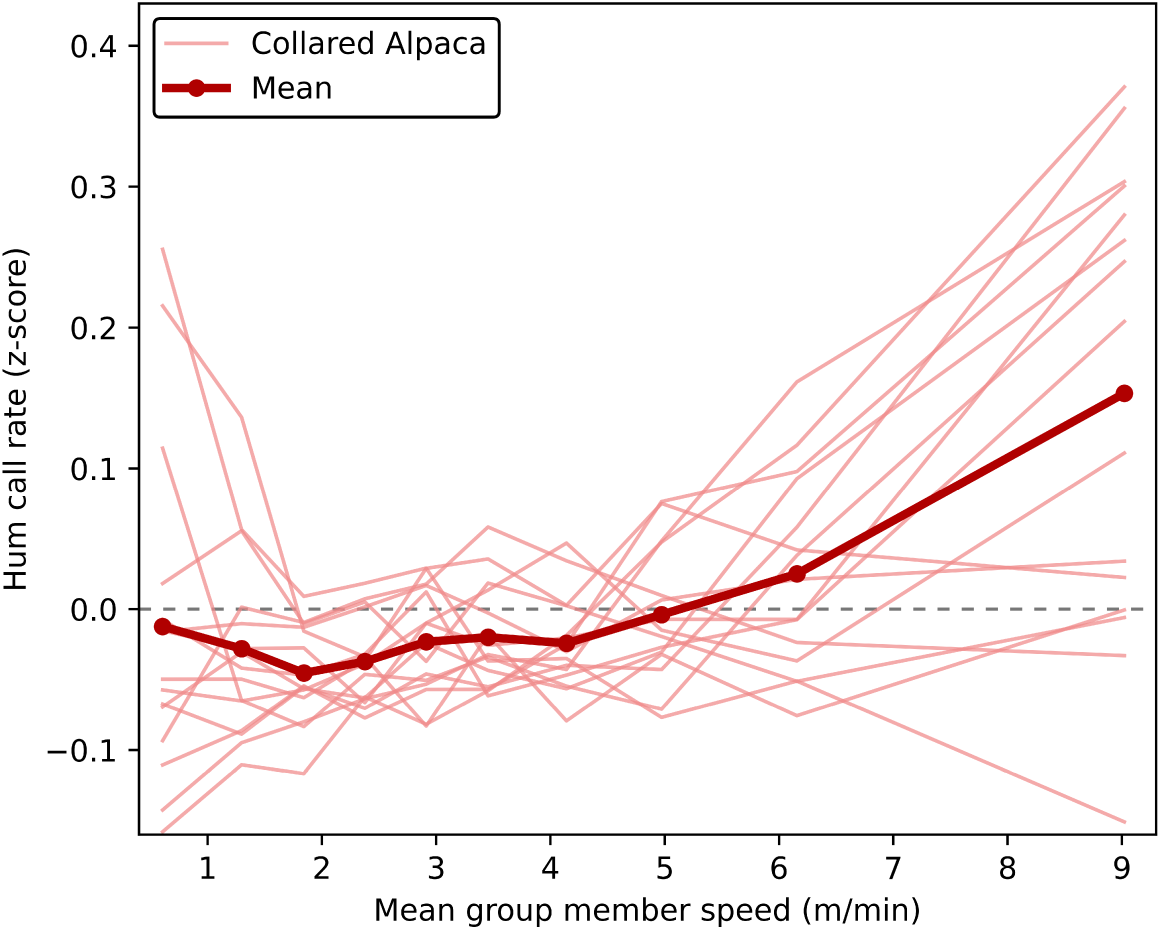
Relationship between *mean group member speed* and standardised hum call rate under the primary 95th-percentile attenuation-calibrated amplitude threshold. Hum call rate was defined as the number of retained hum occurrences per 60-s window. Hum call rate was standardised within focal alpaca. Thin red lines show focal-alpaca means across equal-frequency bins of *mean group member speed*, and the thick red line and points show the mean across focal alpaca. *Mean group member speed* is expressed in m/min. The dashed horizontal line represents the overall mean hum call rate, irrespective of *mean group member speed*.

**Table 4.** Fixed-effect estimates from the primary H2 (Contact Maintenance) negative-binomial type 1 (NB1) model, testing whether hum production was more frequent as relatively compact groups began to spread. The response was *hum count* in 60-s windows, using detections retained above the 95th-percentile amplitude threshold. Predictors were *current spread*, *spread change*, their interaction, *mean group member speed* and *centroid displacement*. Incidence-rate ratios (IRRs) and 95% confidence intervals describe associations with the expected hum count. Continuous predictors were standardised to one standard deviation before model fitting. For *current spread* and *spread change*, the individual IRRs describe their associations when the other interacting predictor is at its mean. *P*-values are from coefficient-level Wald *z*-tests.

| Predictor | IRR | 95% CI | $z$ | $P$ |
| --- | --- | --- | --- | --- |
| Current spread | 0.874 | 0.632–1.209 | -0.813 | 0.416 |
| Spread change | 1.073 | 0.987–1.166 | 1.162 | 0.096 |
| Mean group member speed | 1.179 | 1.119–1.243 | 6.196 | <0.001 |
| Centroid displacement | 1.039 | 0.964–1.121 | 1.001 | 0.317 |
| Current spread × spread change | 0.965 | 0.904–1.029 | -1.082 | 0.279 |

The NB1 interaction model results were consistent when the 90th and 97.5th percentile amplitude thresholds were used instead. The *current-spread × spread-change* interaction remained unsupported at both alternative thresholds (90th percentile: IRR = 0.976, 95% CI: 0.928–1.026, likelihood-ratio test P = 0.363; 97.5th percentile: IRR = 0.960, 95% CI: 0.894–1.030, likelihood-ratio test P = 0.242), while more hums were produced when the collared alpaca were moving faster at both thresholds (Supplementary Fig. S2).

## DISCUSSION

Biologging can generate acoustic and movement data across long recording periods that are difficult to obtain by direct manual observation, but the resulting volume of data can itself become a bottleneck in obtaining useful results, because calls must be extracted before they can be related to behaviour. Here, we addressed this by using automated call detection across continuous recordings and linking detected alpaca hums to GPS-derived spatial data to investigate whether humming presence and frequency was related to group movement and spacing. Our curated dataset allowed us to test two complementary hypotheses at the individual and group level. Hum production preceded alpaca coming closer together, and more hums were produced when the animals were moving faster. Together, these findings l ink h umming to both movement and closer spatial contact, supporting a cohesion-related function of this call type.

At the individual level, focal alpaca tended to be closer to the group centre after producing a hum than during similar moments without a hum. This held true after accounting for their initial distance from the group and whether they were already approaching or moving away from it, indicating that the observed change following a hum was not simply a continuation of the preceding movement pattern. However, because some focal alpaca had already begun moving closer to the group before humming, we cannot conclude that hums initiated this movement, but they clearly accompanied it. Taken together, these findings are consistent with a role for humming in maintaining or restoring social contact as alpacas adjust their positions within the group.

The broader spatial context also mattered. Alpaca that started farther from the group tended to come closer more strongly. Whereas this tendency was weaker when the group as a whole was already more dispersed. By contrast, whether the group had recently been spreading out or contracting did not predict how the focal alpaca’s distance to the group changed next. When we looked more closely at what happened after hum production, the focal alpaca and the group usually came closer together, rather than simply moving apart less. This closer spacing arose in several ways: the focal alpaca moved toward the group, the group moved toward the focal alpaca, or both moved toward one another. Together, these results show that humming was linked to closer contact within a broader pattern of how alpaca adjusted their positions relative to the group, rather than through one fixed movement response. This pattern suggests that humming accompanies the maintenance or restoration of social contact. Familiar alpaca have been observed grazing and resting together, and quiet vocalisations have been described as helping individuals maintain contact (Kapustka and Budzyńska 2018). Separation is also stressful for alpaca; for example, separation during weaning is followed by a temporary increase in cortisol, although this combines maternal separation with a change in environment (Bravo et al. 2001). Our findings extend these observations by showing that humming is associated with closer spacing during routine group activity, supporting a possible role in maintaining social contact before individuals become substantially separated. Mammalian social organisation varies along several dimensions; groups with stable membership can temporarily separate into smaller subgroups while retaining the same members (Prox and Farine 2020). Contact vocalisations may therefore be useful whenever group members become separated in space, whether this occurs in a highly fluid fission–fusion society or within an otherwise stable group. Irrespective of the social organisation in alpaca, which has not yet been characterised, our results provide evidence that the hums maintain or re-establish spatial contact.

Alpaca hums also contain strong individual identity cues (Wierucka and Leu 2024). These cues do not by themselves demonstrate that alpaca recognise callers by voice, but they provide acoustic information that could allow receivers to distinguish between individuals. Such information could be useful during routine group activities in which animals remain socially associated while changing position. Kapustka and Budzyńska (2018), for example, observed familiar alpaca grazing together and described quiet vocalisations as helping them maintain contact. We did not record behavioural state and therefore cannot determine whether the hums in our study occurred during grazing. More generally, however, everyday activities in which group members change their relative positions provide a plausible context for repeated contact maintenance. This interpretation also connects with the group-level result: hums were more frequent when the collared alpaca were moving faster, suggesting that humming may be particularly relevant while group members are actively repositioning.

The increase in humming when the collared alpaca were moving faster was not tied to the specific spacing pattern predicted under H2. Hums were not specifically more frequent as a relatively compact group began to spread, so the group-level analysis did not support this particular prediction for contact maintenance. Instead, the clearest group-level result was that humming was associated with movement. This is biologically relevant because movement can make cohesion more difficult to maintain. In sheep, for example, groups were more spread out when only some individuals were active, whereas groups whose members were active together remained more cohesive, showing that activity can influence how a herd stays together (Michelena et al. 2008). This also connects the group-level and individual-level results. At the group level, hums were more frequent when the alpaca were moving faster, whereas at the individual level, spacing shifted toward closer contact after a hum. Faster movement did not itself explain the individual-level change: the focal alpaca’s speed before the hum was not related to what happened afterwards. Rather, the two results are consistent with humming occurring while alpaca are moving and adjusting their positions relative to one another. Routine grazing provides one plausible context for this. Alpaca spend much of the day grazing, and individuals can change their relative positions while feeding and remaining part of the group (Kapustka and Budzyńska 2018). Humming may therefore help maintain or restore contact during these commonly occurring movements. Future research could combine acoustic and movement recordings with behavioural observations to test whether humming occurs during grazing and how it relates to changes in group spacing during feeding.

Our interpretation that humming may help alpaca maintain social contact during movement is consistent with findings in foraging meerkats. These animals move continuously while foraging and frequently produce soft, short-range contact calls, known as close calls (Gall and Manser 2017). Experimentally manipulating the spatial distribution of these calls caused meerkat groups to elongate and split into subgroups (Gall and Manser 2017). In meerkats, close calls are therefore linked directly to movement during foraging, which is the aspect most similar to our result regarding *mean group member speed*. Close-call exchanges also become more frequent with an increasing number of nearby neighbours, potentially providing repeated information about neighbours’ locations and identities (Demartsev et al. 2024). In a complementary playback experiment, meerkats used individually distinctive close calls to locate a socially important group member and changed their response according to their recent social relationship with that individual (Reber et al. 2013). Future research should investigate whether individual alpaca adjust their movements in response to hums and whether these responses depend on the caller’s identity, location or social relationship to the receiver. Simultaneous acoustic and GPS recordings from multiple collars could help identify likely callers and potential receivers, while playback experiments could test responses to calls from different individuals or apparent locations.

Our design allowed us to collect simultaneous audio and movement data from multiple freely moving alpacas, while requiring us to account for incomplete herd coverage and uncertainty in caller identity. We therefore interpreted the spatial measures as describing the collared subgroup rather than the entire herd. To reduce the likelihood of including hums produced by nearby uncollared alpacas, we applied amplitude thresholds derived from a separate sound-attenuation experiment and repeated the analyses using stricter thresholds. Although GPS-network studies show that the effects of missing individuals vary with the spatial metric and sampling fraction (Silk et al. 2015; Kaur et al. 2024, 2025), our main findings were robust to these sources of uncertainty. Applying stricter amplitude filters produced the same group-level conclusion: humming was more frequent when collared alpacas were moving faster, but was not associated with greater group dispersion or the onset of group spreading. At the individual level, the shift towards closer spacing after humming was consistent across all time intervals examined and remained after unusually distant alpacas were excluded. Together, these sensitivity analyses show that partial collaring can provide robust and biologically meaningful insights into vocal and spatial behaviour, even when complete herd coverage is not feasible. Future work could assess the effects of sampling fraction through simulation studies or by collecting full-coverage datasets and systematically removing individuals to test how well patterns derived from partial samples represent whole-group dynamics.

Our study shows how automated call detection, followed by curation of the initial detections, can make extensive acoustic recordings usable for behavioural research. Across 2,011 h of continuous recordings, this approach yielded 28,561 curated hum occurrences that could be linked to GPS-derived measures of group movement and spacing. Detecting candidate calls automatically made processing this volume of recordings feasible, while subsequent curation and duplicate removal reduced false detections and duplicate recordings before analysis. In alpaca, the associations between hum production and spatial arrangement seem to provide a reliable basis for investigating the possible role of this vocalisation in maintaining social contact. Furthermore, identifying repeated associations between vocalisations and behaviour narrows the plausible functions of calls, providing a basis for designing future targeted observational and experimental studies. Combining scalable acoustic processing with synchronised movement data therefore provides a powerful approach for investigating vocal communication in species whose behaviour and call functions are poorly understood.

## ACKNOWLEDGEMENTS

We thank Yunus Emre Atalay for conducting the initial annotations used to train the CNN detector. We also thank Dr. Roger Mundry for insight and advice on the linear mixed-effects models. We thank Prof. Alexander Ecker for supervision and input. We thank the KISSKI project for granting computing time (the calculations for this research were conducted with computing resources under the project kisski-alpaca-2). ROHR was supported by a Summer Research Scholarship from Adelaide University.

## DECLARATION OF GENERATIVE AI USE

OpenAI ChatGPT (GPT-5.6 Sol, high reasoning; OpenAI 2026) was used for language improvement, including grammar and sentence structure. Anthropic Claude Code (Claude Opus 4.8, high effort; A nthropic 2 026) was u sed for coding assistance during development and debugging, and for basic assistance with the LATEX project. The authors retain full responsibility for the final manuscript and analyses.

## DATA AND CODE AVAILABILITY

The code for automated hum detection and subsequent candidate curation is publicly available at https://github.com/darizae/alpaca-pipelines.

