## Supplementary Materials for "Evidence for a cohesion-related function of alpaca hums: Linking automated call detection with GPS data"

**Table S1.** Selected duplicate-removal thresholds and grid-search diagnostics. Candidate pairs linked detections from different collars whose intervals overlapped by at least the selected fraction of the shorter detection and whose collar positions were within the selected distance. Acoustic agreement was assessed across seven descriptors: Fundamental Frequency ($F_0$), Duration 90%, Peak Frequency, Frequency 5%, Frequency 95%, Bandwidth 90% and Aggregate Entropy (Charif et al. 2010). For provisional groups of three or more detections, descriptors were standardised, pairwise Euclidean acoustic distances were calculated and the selected group threshold was applied to the candidate-distance distribution. Detections connected by accepted similarity edges were treated as one occurrence, and one representative per connected component was retained by prioritising signal-to-noise ratio, then root-mean-square amplitude and detector score. The selected configuration was chosen from the 500-setting grid on the basis of acoustic coherence and stability of duplicate decisions and removal counts rather than maximum removal.

| Quantity | Selected value or diagnostic |
|---|---|
| Threshold combinations evaluated | 500 |
| Minimum temporal overlap of shorter detection | 0.05 |
| Maximum collar distance | 6 m |
| Maximum pairwise relative acoustic difference | 0.10 |
| Provisional-group acoustic-distance threshold | q90 |
| Candidate edges between AudioMoths | 903 |
| Accepted acoustic-similarity edges | 326 |
| Multi-detection events | 116 |
| Maximum connected-component size | 8 |
| Local decision stability | 0.958 |
| Removal-count stability | 0.977 |
| Overall robustness | 0.978 |

**Table S2.** A priori interpretation of the H1 hum-presence coefficient $\beta_{\text{hum}}$. The coefficient compares mean change in focal-to-centroid distance ($\Delta D_w$ ) between hum occurrences and matched controls after adjustment for the other model predictors. Its sign therefore describes a difference between conditions and does not, by itself, determine whether distance decreased after every hum.

| **Hum-presence coefficient** $\beta_{\text{hum}}$ | **Meaning for hum versus control distance change** | **A priori H1 interpretation** |
|---|---|---|
| Negative | Mean $\Delta D_w$ is more negative after hums than controls. This can arise because focal-to-centroid distance decreases after hums, or because distance increases in both conditions but increases less after hums. | Predicted H1 direction; consistent with a cohesion-oriented contact-reinforcement effect. The descriptive breakdown of matched pairs in the primary 60-s H1 analysis distinguishes these two possibilities. |
| Around zero | Mean $\Delta D_w$ does not differ detectably between hum and control occurrences. | No detectable support for the predicted H1 effect at that interval length. |
| Positive | Mean $\Delta D_w$ is more positive after hums than controls, for example through greater post-hum expansion or less post-hum contraction. | Inconsistent with the predicted cohesion-oriented H1 effect. |

**Table S3. A priori interpretation of the H2 model terms.** Incidence-rate ratios (IRR) describe the multiplicative change in fitted hum count for a one-SD increase in a predictor, with the other predictors held constant. *Current spread* increases with dispersion and *spread change* is positive during expansion, so the conditional pattern predicted under H2 corresponds to an interaction IRR below one.

| **Model term and question** | **Result expected under H2 (IRR)** | **Interpretation if H2-consistent** | **Interpretation if H2-inconsistent** |
|---|---|---|---|
| *Current spread* — Are hums more frequent when the group is compact? | A one-SD increase in spread is associated with fewer expected hums, at average *spread change* (IRR $< 1$). | Humming is associated with compact, cohesive group states and is less frequent when dispersion is high. | Humming is unrelated to compactness, or is more frequent in dispersed groups. |
| *Spread change* — Are hums more frequent while spread is increasing? | A one-SD increase in *spread change* is associated with more expected hums, at average *spread* (IRR $>$ 1). | Humming is associated with the group expanding and cohesion decreasing, and is less common during contraction. | Humming is unrelated to expansion, or is more frequent during contraction. |
| *Mean group member speed* — Are hums more frequent when *mean group member speed* is higher? | A one-SD increase in *mean group member speed* is associated with more expected hums (IRR $> 1$). | Humming is associated with greater movement activity among group members. | No specific prediction. |
| *Centroid displacement* — Are hums more frequent when *centroid displacement* is greater? | A one-SD increase in *centroid displacement* is associated with more expected hums (IRR $>$ 1). | Humming is associated with greater displacement of the group centre. | Humming is not associated with displacement of the group centre; this does not independently reject H2. |
| *Current spread* × *spread change* — Are hums more frequent when the group is compact yet expanding? | The interaction IRR is below one, so the positive association between *spread change* and hum count is stronger at low *current spread*. | The effects of *spread* and *spread change* depend on each other: humming in compact groups is more frequent when dispersion is increasing than when it is not, supporting contact maintenance. | The effects of *spread* and *spread change* are independent, giving no support for contact maintenance. |

**Table S4. Detailed evaluation of the automated corpus-construction layers.** Evaluation designs, sample sizes and performance measures are reported for the CNN candidate generator, the random-forest and logistic-regression curation layers, and the threshold search used for duplicate removal.

| **Layer** | **Evaluation material** | **Evaluation design** | **Results** |
|---|---|---|---|
| CNN candidate generation | 4,325 labelled excerpts: 1,730 hums and 2,595 noise examples (3,058 training; 570 validation; 697 test) | Training, validation and test partitions; checkpoint selected by validation accuracy | Best validation accuracy 0.960 at epoch 32; test accuracy 0.968; F1 0.964; precision 0.970; recall 0.958 |
| Random-forest candidate curation | 3,998 reviewed candidates from 29 recordings: 1,730 hums and 2,268 non-target detections | Grouped 75:25 partition; 150 model and decision-threshold combinations | Reviewed candidate hum share 43.3%; accuracy 0.805; F1 0.749; precision 0.747; recall 0.752; ROC AUC 0.875; TN 374, FP 72, FN 70, TP 212 |
| Logistic contamination filtering | 1,390 reviewed detections from 185 source WAVs: 1,105 hums, 275 human-speech examples and 10 rumbles | Five-fold cross-validation grouped by source WAV | Balanced accuracy 0.801; accuracy 0.817; ROC AUC 0.896; hum precision 0.935; hum recall 0.827; noise recall 0.775; TN 221, FP 64, FN 191, TP 914 |
| Duplicate removal | 28,814 detections; 500 threshold configurations evaluated | Selection based on local decision stability, acoustic coherence and removal-count stability; selected thresholds: 5% overlap, 6 m GPS distance, 0.10 relative acoustic difference and 90th-percentile group acoustic distance | Robustness 0.978; local decision stability 0.958; removal-count stability 0.977; 28,561 retained occurrences |

**Table S5. Directional cases among matched hum-control pairs in the primary 60-s H1 analysis.** Cases are defined by the signs of the change in focal-to-centroid distance($\Delta D_{60}$) for hum presence and, when both changes have the same sign, by which condition has the larger change. The final column gives the percentage of classified matched hum–control pairs assigned to each case.

| Directional case | Mean hum ΔD (cm) | Mean control ΔD (cm) | Proportion (%) |
|---|---|---|---|
| Greater increase following hum than matched control | 384 | 95 | 13.88 |
| Smaller increase following hum than matched control | 122 | 347 | 9.93 |
| Decrease following hum; increase at matched control | -445 | 238 | 26.55 |
| Greater decrease following hum than matched control | -624 | -132 | 16.35 |
| Smaller decrease following hum than matched control | -158 | -418 | 10.92 |
| Increase following hum; decrease at matched control | 261 | -163 | 22.37 |

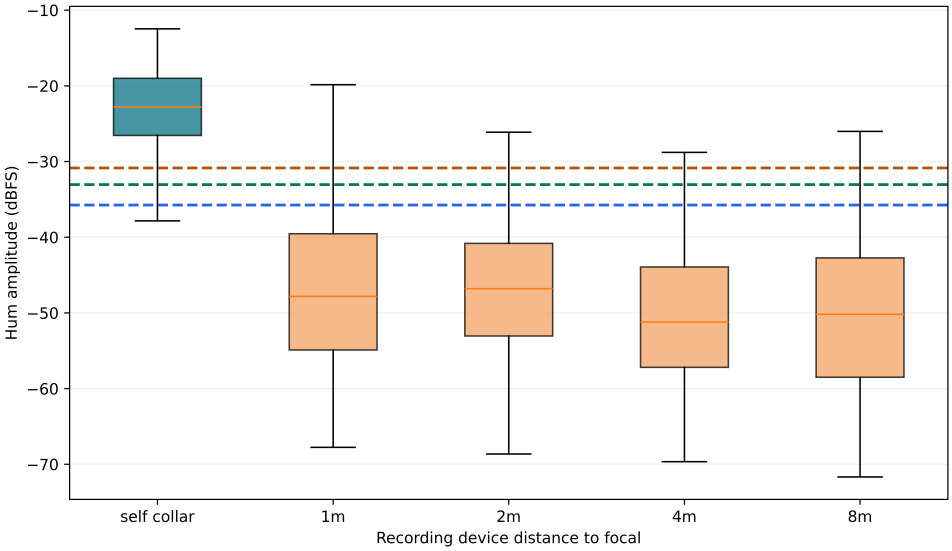


**Figure S1.** Calibration of the amplitude thresholds used in H2 (Contact Maintenance) to reduce the inclusion of hums produced by nearby alpaca rather than the collar wearer. Boxplots show the 95th-percentile hum-band root-mean-square (RMS) amplitude, a measure of recorded hum signal strength, for hums recorded on the caller's own collar and simultaneously by stationary AudioMoths positioned 1, 2, 4 and 8 m from the caller. Horizontal dashed lines indicate thresholds derived from the pooled stationary-AudioMoth amplitude distribution: the 90th percentile (-35.75 dBFS), 95th percentile (-33.06 dBFS) and 97.5th percentile (-30.85 dBFS), in order of increasing amplitude. The 95th-percentile threshold (-33.06 dBFS) was used for the primary H2 statistical analyses, while the other two thresholds were used for sensitivity analyses.

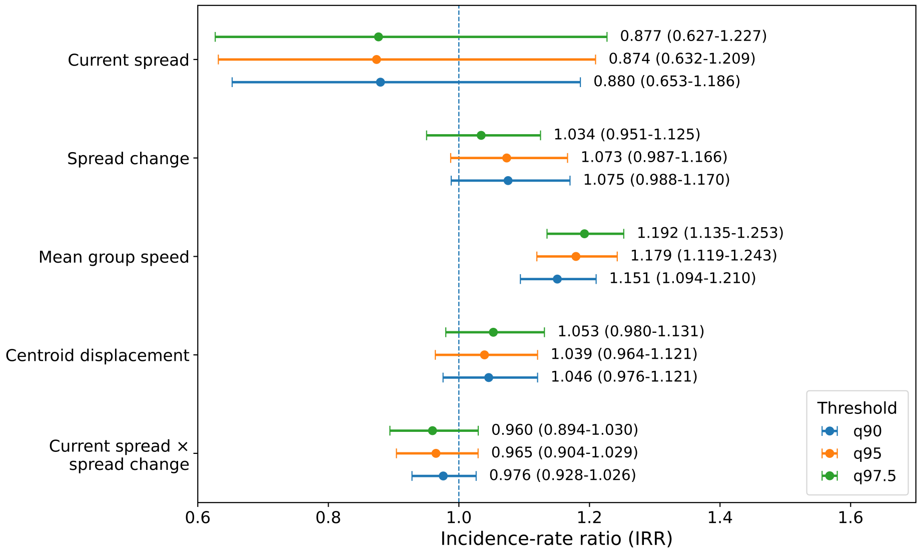


**Figure S2.** Fixed-effect incidence-rate ratios (IRRs) from the NB1 model with the *current-spread* × *spread-change* interaction across the 90th, 95th (primary), and 97.5th percentile attenuation-calibrated amplitude thresholds. Points show IRRs and horizontal bars show 95% Wald confidence intervals for a one-SD increase in each predictor. The vertical dashed line marks an IRR of one. Across all three thresholds, *mean group member speed* was positively associated with hum production, whereas the *current-spread* × *spread-change* interaction was not supported.
